# Runx1 Shapes the Chromatin Landscape Via a Cascade of Direct and Indirect Targets

**DOI:** 10.1101/2020.09.25.313767

**Authors:** Matthew R. Hass, Daniel Brisette, Sreeja Parameswaran, Mario Pujato, Omer Donmez, Leah C. Kottyan, Matthew T. Weirauch, Raphael Kopan

## Abstract

Runt-related transcription factor 1 (Runx1) can act as both an activator and a repressor. Here we show that CRISPR-mediated deletion of *Runx1* in an embryonic kidney-derived cell (mK4) results in large-scale genome-wide changes to chromatin accessibility and gene expression. Open chromatin regions near down-regulated loci are enriched for Runx sites, remain bound by Runx2, but lose chromatin accessibility and expression in *Runx1* knockout cells. Unexpectedly, regions near upregulated genes are depleted of Runx sites and are instead enriched for Zeb transcription factor binding sites. Re-expressing Zeb2 in *Runx1* knockout cells restores suppression. These data confirm that Runx1 activity is uniquely needed to maintain open chromatin at many loci, and demonstrate that genome-scale derepression is an indirect consequence of losing Runx1-dependent Zeb expression.

## Introduction

Transcription factors (TFs) play fundamental biological roles by controlling gene expression, the first step in translating genomic DNA sequence into function. Mammalian genomes encode over 1,000 TFs, which precisely control gene expression through complex combinatorial interactions and transcriptional cascades (Lambert et al., 2018). To achieve this precision, TFs use a wide range of mechanisms, including initiating the activation or repression of gene expression through the recruitment of co-factors, initiating new chromatin looping interactions between enhancers and target promoters, and altering the chromatin landscape through the repositioning of nucleosomes (Lee and Young, 2013). Achieving an understanding of the mechanisms underlying the control of gene expression is thus an enduring and fundamental goal of molecular biology.

Runx/Runt TF family members recognize a characteristic TGTGGT DNA-binding motif, and are present across all metazoans (Rennert et al., 2003). Members of the Runx family play important roles in development and disease (Ito et al., 2015; Mevel et al., 2019), notably during hematopoiesis (de Bruijn and Dzierzak, 2017; Seo and Taniuchi, 2020), skin development (Glotzer et al., 2008; Hoi et al., 2010; Osorio et al., 2008), and ossification (Komori, 2018; Mevel et al., 2019; Sierra et al., 2004; Zhang et al., 2008a). Runx proteins can act as repressors, by recruiting the Groucho/TLE proteins via a C-terminal tetrapeptide WRPY, or as activators, by heterodimerizing with Core binding factor (CBF)ß and recruiting cell context-specific activators. In several developmental contexts, Runx proteins collaborate with Notch, at times facilitating Notch activity (Giambra et al., 2012; Terriente-Felix et al., 2013), and at others acting downstream of Notch (Kueh et al., 2016). The role that Runx1 plays in establishing chromatin accessibility has been studied in some detail within the hematopoietic system (Lichtinger et al., 2010), where it acts as a pioneer protein. However, how Runx proteins influence gene regulatory networks in different cellular contexts remains to be elucidated.

Previously, we found that Runx binding sites were enriched near Notch-bound enhancers in a diploid kidney metanephric mesenchymal cell line, mK4 (Hass et al., 2015). Two of the three Runx orthologs, Runx1 and Runx2, are expressed in the kidney-derived mK4 cells (Valerius et al., 2002), facilitating detailed molecular comparison of Runx1 versus Runx2 functions in regulating gene expression and their integration with multiple signaling pathways. Such analyses are further aided in mK4 cells by the normal karyotype, the ease of CRISPR-mediated genetic manipulation, and by short replication times, providing sufficient material for a variety of genomic assays.

In this study, we show that Runx1 plays an important role in regulating chromatin accessibility at many genomic loci in mK4 cells. In the absence of Runx1, Runx2 bound most of the Runx1-bound chromatin but could not maintain Runx1-dependent accessibility or gene expression. As Runx1 can repress expression of some genes, we anticipated these to be re-expressed in Runx1KO cells; however, we were surprised to discover that accessible chromatin near loci expressed only after Runx1 deletion were instead enriched for Zeb sites and depleted of Runx sites, suggesting indirect involvement of Runx1 in their regulation. Further investigation revealed that repression at multiple loci throughout the genome is mediated by two Runx1-dependent targets, Zeb1 and Zeb2.

Restoring Zeb2-expression in Runx1KO cells restored repression of target genes. Thus, the direct impact of Runx1 on chromatin in mK4 cells is mediated primarily through its pioneer and transcriptional activator function, rather than through its repressor function. Collectively, these data reveal an important role for Runx TFs in the maintenance of the chromatin landscape and provide mechanistic insight into how Runx and Zeb TFs interactively control gene expression in the kidney.

## Results

### Generation and characterization of Runx1 knockout cells

To generate cells lacking Runx1 activity, we targeted Runx1 with two gRNAs flanking exon 3, which contains the start codon of the transcript expressed in mK4 cells and encodes part of the Runt DNA binding domain (Figure 1A). Multiple clones grew after selection with puromycin showing deletion of the targeted exon 3 (Figure 1A) by PCR genotyping and loss of Runx1 protein as confirmed by Western blot (Figure 1B). The expression of Runx2 in these cells remained unchanged (Figure 1B; Figure 1-Supplemental Figure 1). Thus, any functional differences between Runx1KO cells and the parental cell line (control) would indicate potential Runx1-specific roles that cannot be compensated for by Runx2.

**Figure 1:**
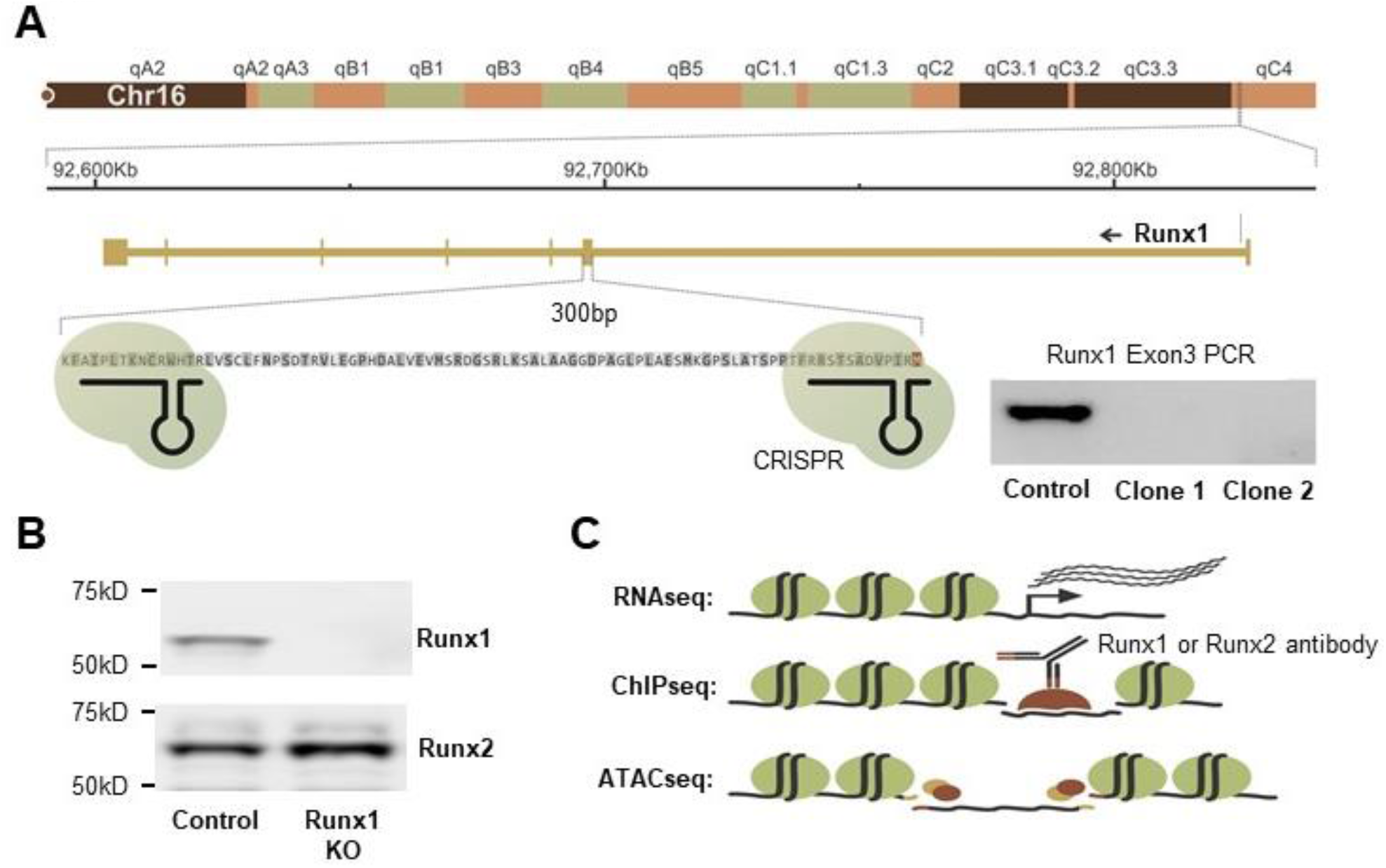
Generation and Characterization of Runx1KO Cells. A) Diagram of the Runx1 exon 3 region targeted for deletion using CRISPR-Cas9 and confirmation of deletion by PCR. B) Western blot showing that Runx1KO cells lack Runx1 protein but contain Runx2. C) Schematic of genomic analyses utilized to characterize Runx1KO cells.

To determine the impact of Runx1-deficiency, we performed multiple genomic assays comparing Runx1KO to control mK4 cells. Specifically, we analyzed gene expression through RNA-seq, identified genomic locations bound by Runx1 and Runx2 through ChIP-seq, and mapped chromatin architecture by ATAC-seq (Figure 1C). The RNA-seq, ChIP-seq, and ATAC-seq experiments were all performed in biological triplicates to enable statistical analyses for the identification of significant differences between control and Runx1KO cells.

### Runx1-deficiency induces dramatic changes in gene expression

RNA-seq analysis identified thousands of genes that were significantly altered in Runx1KO cells relative to control cells (Figure 2A). The replicates were highly consistent and revealed 1,705 upregulated and 1,182 down-regulated transcripts in the Runx1KO cells (fold change > 2-fold, FDR < 0.05 across replicates) (Supplemental Table 1). GO term analysis of the upregulated genes in Runx1KO cells identified enrichment for the biological process of antigen processing and presentation (5.7 fold enriched, p-value 0.03, Supplemental Table 2), consistent with the critical role that Runx1 plays in the immune system and with observations in human patients with Runx1 mutations (Awad et al., 2018). Consistent with previous studies, Runx1KO cell downregulated genes were enriched for the TGF-beta receptor signaling pathway (23.5 fold enriched, p-value 0.0012, Supplemental Table 2) (Zhou et al., 2018). The widespread changes in gene expression caused by Runx1-deficiency are consistent with previous observations of non-redundancy with Runx2, as seen in other cellular contexts (Mevel et al., 2019). Thus, Runx1 plays a critical role in controlling the transcriptome within mK4 cells in a manner that cannot be compensated for by Runx2. We next examined if differences between Runx1 and Runx2 effects on gene expression might be due to differences in DNA binding preferences or differences in genomic binding locations.

**Figure 2:**
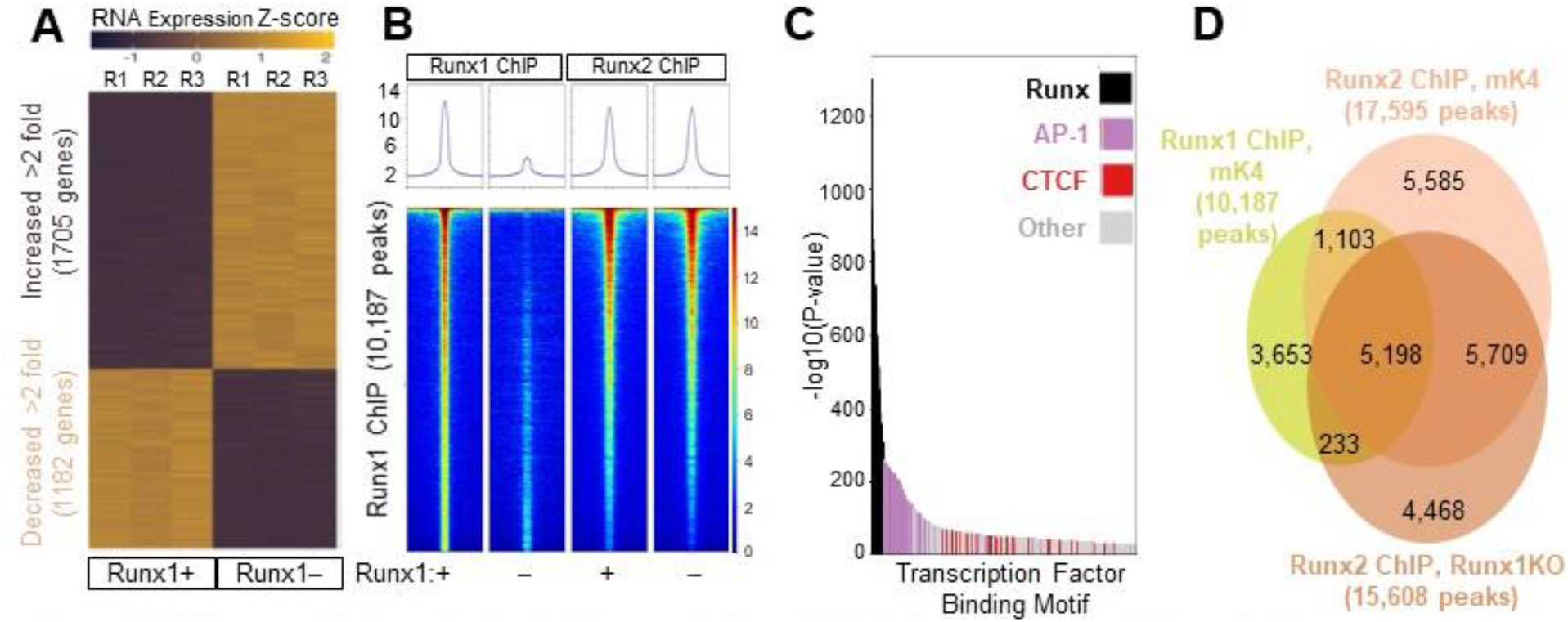
Widespread Transcriptional Changes in Runx1KO Cells Despite Runx2 Largely Occupying the Same Regions as Runx1. A) Heatmap of RNA-seq triplicates showing 1,705 upregulated and 1,182 downregulated genes (over 2 fold) in Runx1KO cells compared to control mK4 cells. B) Heatmap of ChIP-seq reads mapping to Runx1 peaks from Runx1 ChIP or Runx2 ChIP in control versus Runx1KO cells. C) Graph displaying −log10 p-values of motif enrichment, revealing that Runx1 motifs are the most highly enriched motifs in the Runx1 ChIP peaks. D) Venn diagram showing strong overlap of Runx1 and Runx2 ChIP peaks.

### Runx1 and Runx2 bind near genes that are down-regulated in Runx1KO cells

To identify genomic loci occupied by Runx proteins, we performed Runx1 and Runx2 ChIP-seq. The specificity of the Runx1 antibody used for ChIP was determined by performing the experiment in control and Runx1KO cells. ChIP-seq in control cells produced 10,187 peaks that were highly reproducible between replicates but not present in Runx1KO cells, as depicted in heatmaps of the Runx1 ChIP-seq reads mapped to peaks in control and Runx1KO cells (Figure 2B; Figure 2-Supplemental Figure 1A). HOMER transcription factor binding site motif enrichment analysis using mouse motifs from the Cis-BP database (Lambert et al., 2019) identified Runx motifs as most highly enriched in the control cell dataset (p-value 1 × 10^−1298^) (Figure 2C and Supplemental Table 3). These data, combined with the limited number of peaks and relative lack of Runx motif enrichment in the Runx1KO cells, confirm the specificity of the antibody and that the identified genomic regions are bound by Runx1. The second most enriched class of motifs were for the AP-1 family. Runx1 genomic binding has previously been shown to be co-enriched with members of this family (Pencovich et al., 2011). Co-association with AP-1 suggested that many of the Runx1-bound regions are likely enhancers, given the known association of AP-1 sites with enhancers in most cell types (Andersson et al., 2014).

To further determine whether these Runx1 bound regions were involved in transcriptional regulation, we assigned genes to the Runx1 ChIP peaks using the GREAT annotation tool (McLean et al., 2010) and compared the genes near immunoprecipitated chromatin to the genes that exhibiting expression changes of over 2-fold in Runx1KO cells (Supplemental Table 1). This analysis showed that 45% (526/1182) of downregulated genes in Runx1KO cells had a Runx1 ChIP-seq peak in their vicinity, a 1.85 fold enrichment over what was expected by chance (hypergeometric p-value 4.60e × 10^−55^). In contrast, only 30% (511/1705) of upregulated genes had a nearby immunoprecipitated peak (a 1.24 fold enrichment). These results are consistent with the loss of expression in Runx1KO cells of genes predicted to be activated by Runx1 but are less consistent with a model in which upregulated genes were repressed directly by Runx1.

The failure of Runx2 to regulate the same genes as Runx1, as reflected in the RNA-seq data (Figure 2A), might reflect physical occlusion of Runx2 binding by Runx1. Alternatively, certain regions of the genome might only be occupied by Runx1, and not Runx2, due to differences in DNA binding preferences or protein interaction partners. Finally, Runx1 and Runx2 might occupy the same loci, but Runx2 might have different effects on gene expression compared to Runx1. To further investigate these possibilities, we performed Runx2 ChIP-seq in both control and Runx1KO cells. The Runx2 ChIP identified combined sets of 17,595 peaks present in control cells and 15,608 peaks present in Runx1KO cells, with the Runx motif strongly enriched in both cell types (p-value 1 × 10^−2244^ and 1 × 10^−2363^, respectively (Supplemental Table 3)). Comparisons between the chromatin bound by Runx1 and Runx2 revealed remarkable overlap of the peaks in control cells, and retention of Runx2 ChIP signal at Runx1 peaks in Runx1KO cells (Figures 2B, 2D, and Figure 2-Supplemental Figure 1A). Regulatory Element Locus Intersection (RELI) analyses (Harley et al., 2018) confirmed the highly significant agreement between the Runx1 (mk4), Runx2 (mk4), and Runx2 (Runx1 KO) ChIP-seq datasets (control cell 194.46 fold enriched, p-value 2.0 × 10^−219^; Runx1KO cell 177.85 fold enriched, p-value 2.32 × 10^−219^) (Figure 2-Supplemental Figure 1B and Supplemental Table 4). These results suggest that the regulatory regions near downregulated genes in Runx1KO cells retain Runx2 binding, which evidently is not sufficient to drive their expression. Further, the ChIP data indicate that Runx1 has transcriptional activator function at a large subset of its target loci. Runx2 can also bind these loci, but this binding is not sufficient to activate the expression of the associated genes

### Dramatic changes in chromatin accessibility drive expression changes in Runx1KO cells

The Runx ChIP data indicated that most regulatory regions remained accessible to Runx2 binding in Runx1KO cells (Figure 2B). To examine chromatin accessibility near down regulated genes in Runx1KO cells, we next performed ATAC-seq experiments. As with the RNA-seq and ChIP-seq data, all ATAC-seq replicates were highly reproducible (Figure 3-Supplemental Figure 1A). The majority of open chromatin regions represented by 37,481 ATAC-seq peaks displayed similar levels of reads between control and Runx1KO cells, henceforth called Runx1-independent ATAC-seq peaks (Figure 3A, intersect in Venn diagram, and heatmap in Figure 3B, left panel). Notably, we also observed substantial and reproducible loss in chromatin accessibility after Runx1 was deleted – 8,741 genomic loci had significantly lower accessibility in Runx1KO cells vs control, which we denote as Runx1-dependent peaks (Figure 3A, left unique area in Venn diagram and heatmap in Figure 3B, middle panel). Interestingly, a similar number of regions (9,427) showed increased accessibility in Runx1KO cells vs control (Runx1KO-induced; Figure 3A right unique area in Venn diagram and heatmap in Figure 3B, right panel). We denote these sites as Runx1KO-induced.

**Figure 3:**
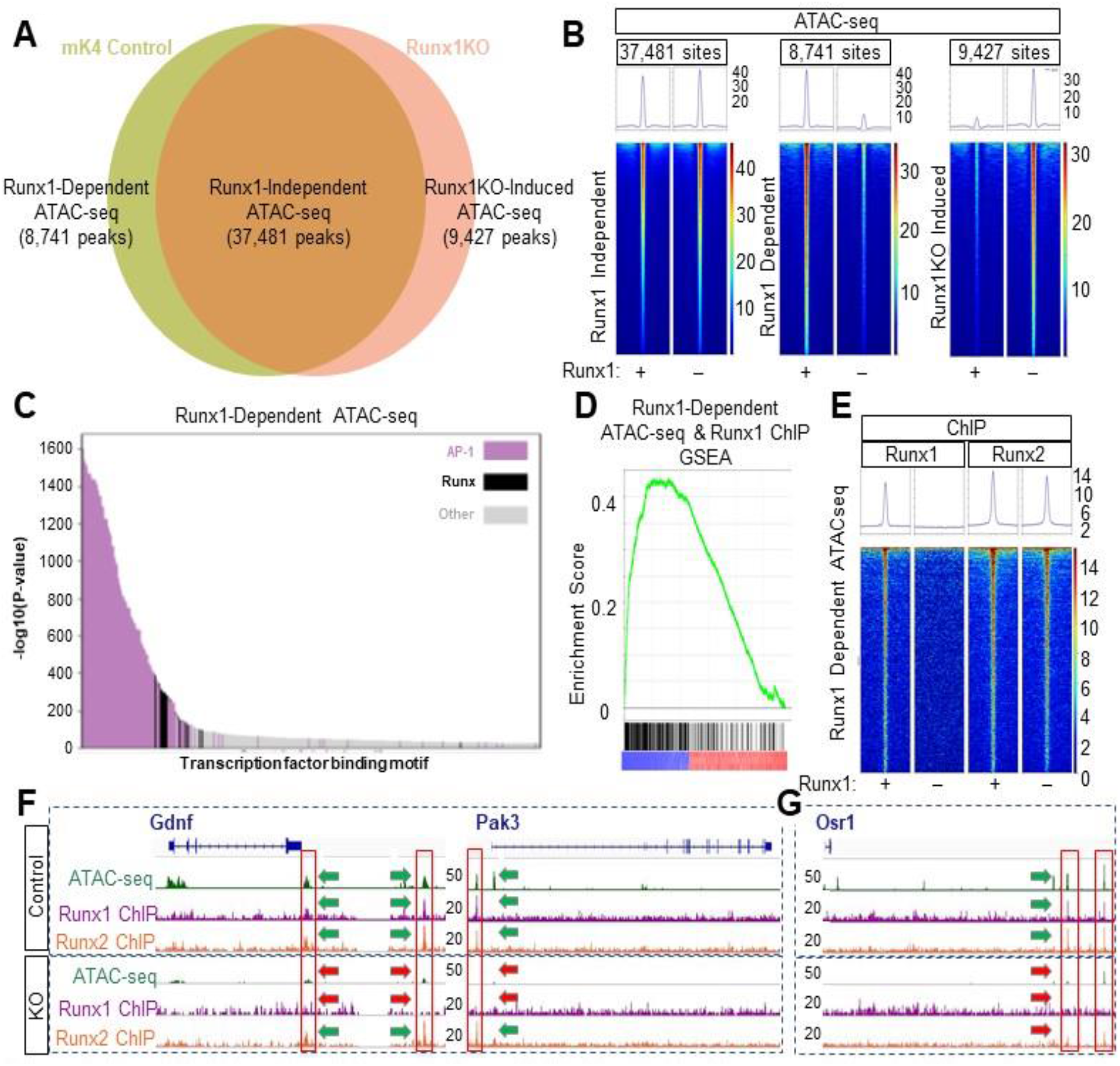
Runx1 Deletion Alters Chromatin Accessibility Despite the Presence of Runx2. A) Venn diagram of ATAC-seq peaks in control and Runx1KO cells showing the number of regions open in both cell lines (Runx1-independent), regions open only in control cells (Runx1-dependent) and regions open only in Runx1KO cells (Runx1KO-induced). B) Heatmaps of the ATAC-seq reads mapping to Runx1-independent, Runx1-dependent, and Runx1KO-induced peaks in the control and Runx1KO cells. C) Graph of the −log10 p-values of motif enrichment, displaying that Runx1-dependent ATAC-seq peaks are strongly enriched for AP-1 and Runx motifs. D) Gene set enrichment analysis showing enrichment of transcriptionally down-regulated genes by Runx1-dependent ATAC-seq peaks that are bound by Runx1. E) Heatmap showing Runx1-Dependent ATAC-seq regions bound in Runx1 and Runx2 ChIP experiments. F) Genomic snapshots of *Gdnf* and *Pak3* genes that are downregulated in Runx1KO cells showing open chromatin regions present in control cells but not Runx1KO cells that are bound by Runx1 and Runx2 and retain Runx2 binding in the Runx1KO cells. G) Genomic snapshot of the Runx1KO downregulated gene *Osr1* that has genomic regions that lose chromatin accessibility in Runx1KO cells, which are bound by Runx1 and Runx2 in control cells, with reduced Runx2 binding in Runx1KO cells.

These changes in chromatin accessibility could reflect Runx1 functioning as an activator at some loci (i.e., by opening or maintaining the accessibility of Runx1-dependent sites) and a repressor at other loci (i.e., by keeping Runx1KO-induced sites inaccessible). If Runx1 is directly acting as both an activator and a repressor in this manner, then both classes would be expected to be enriched for Runx1 motifs. To test this hypothesis, we repeated the analyses described above and again found strong enrichment for AP-1 motifs (p-value 1 × 10^−1601^), implicating Runx1-dependent ATAC-seq regions as likely enhancers (Figure 3C and Supplemental Table 3). The 2nd most enriched motif class in Runx1-dependent enhancers was Runx (p-value 1 × 10^−388^) and as expected, the dataset had highly significant overlap with our Runx1 ChIP data (RELI: 42.60 fold enriched, p-value 8.6 × 10^−215^) (Supplemental Table 4). Additionally, the Runx1-dependent regions may be functionally important in multiple cellular contexts as the RELI analysis also found significant enrichment for Runx1 ChIP sites in AML (10.73 fold enriched, p-value 1.62 × 10^−170^) and HPC-7 cells (9.73 fold enriched, p-value 2.18 × 10^−146^) (Supplemental Table 4). Thus, Runx1 is likely playing an active role in keeping these regions accessible in mK4 cells and potentially also in cancer cell types in which Runx1 is known to play a critical role. We next assigned Runx1-dependent ATAC-seq regions to nearby Runx1-dependent transcripts using the GREAT annotation tool (McLean et al., 2010). Runx1-dependent ATAC-seq regions were enriched 3.11 fold near down-regulated genes in Runx1KO cells (hypergeometric p-value = 1.5 × 10^−197^), indicating that the likely enhancers identified by ChIP and ATAC-seq largely act by maintaining expression of nearby genes.

Comparison of Runx1 ChIP peaks obtained in control cells to the three classes of ATAC-seq peaks shown in Figures 3A and 3B revealed extensive Runx1 binding within both Runx1-dependent peaks and peaks that remain open in the absence of Runx1 (Figure 3-Supplemental Figure 1B), which do not require Runx1 to remain accessible. Interestingly, regions that become inaccessible in Runx1KO cells (Runx1-dependent) that are bound in the Runx1 ChIP show strong enrichment (4.03 fold, hypergeometric p-value 1.01e × 10^−67^) for proximal genes whose expression decreases in Runx1KO cells (Figure 3D). Thus, the combination of ATAC-seq and ChIP data helps to define a set of functional, Runx1-dependent regulatory regions in the mK4 genome and supports the hypothesis that Runx1 pioneer/activator function is maintaining accessible chromatin in mK4 cells.

The Runx1-dependent regions that close in the absence of Runx1 despite the presence of Runx2 expression might do so because those specific regions are bound only by Runx1 and not by Runx2. To test this hypothesis, we compared the Runx2 ChIP-seq reads with the three classes of ATAC-seq peaks shown in Figure 3A. As we observed for Runx1 ChIP, the Runx2 ChIP signal was present at both Runx1-independent and Runx1-dependent ATAC-seq regions, and notably, they are clearly present in Runx1KO cells (Figure 3-Supplemental Figure 1B). However, Runx2 ChIP signal was slightly decreased at Runx1-dependent ATAC-seq sites in Runx1KO cells, likely due to decreased chromatin accessibility (Figure 3E). For example, putative enhancers located near the Runx1KO regulated genes *Gdnf* and *Pak3* are shown in Figure 3F (green arrows). Both *Gdnf* and *Pak3* are members of signaling pathways previously reported to be regulated by Runx (Chen et al., 2006; Ernsberger, 2008; Luo et al., 2007; Park et al., 2012; Rouillard et al., 2016). In Runx1KO cells, they became inaccessible (Figure 3F, red arrows) while still retaining Runx2 binding (Figure 3F, green arrow). Further examples are provided as supplemental data to demonstrate that this pattern of lost ATAC-seq signal but retained Runx2 binding is widespread near genes whose expression is reduced in Runx1KO cells (Figure 3-Supplemental Figure 1D). These data suggest that while Runx2 can bind to closed chromatin like Runx1 (Lichtinger et al., 2010), it cannot make the chromatin accessible to other factors.

Other Runx1-dependent regions display greatly reduced Runx2 binding. For example, two potential enhancers near *Osr1*, a reported Runx target gene (Stock et al., 2004), are open and bound by both Runx1 and Runx2 in control cells (Figure 3G, green arrows), but become inaccessible with limited binding of Runx2 in the Runx1KO cells (Figure 3G, red arrows). We tested the hypothesis that sites that lose Runx2 binding in Runx1KO cells may be enriched for downregulated genes by separating the Runx1-dependent ATAC-seq regions bound by Runx1 into two groups: those sites that had an overlapping Runx2 ChIP peak in Runx1KO cells and those sites that did not (Figure 3-Supplemental Figure 1E). Enrichment analysis on these two classes, performed as above, revealed similar strong enrichment for downregulated genes in the sites immunoprecipitated by Runx2 in Runx1KO cells (4.12 fold enrichment, hypergeometric p-value 1.95e × 10^−61^) as well as the sites that are not immunoprecipitated by Runx2 in Runx1KO cells (3.88 fold enrichment, hypergeometric p-value 1.51 × 10^−13^). These results indicate that loss of both Runx1 and Runx2 binding does not compromise expression more than loss of Runx1 alone. Collectively, the Runx1 and Runx2 ChIP data indicate that Runx1 binds to and actively opens or maintains chromatin accessibility at a large number of loci, many of which are associated with Runx1-responsive genes. Despite remaining bound to these same regions in the absence of Runx1, Runx2 binding is unable to compensate for the lack of Runx1.

### Runx1KO-induced chromatin regions are opened due to the loss of Zeb transcriptional repressors

Our analyses above suggest that Runx1KO-induced ATAC-seq regions require Runx1 to remain inaccessible, but the lack of Runx1 binding to these regions is inconsistent with direct repression by Runx1. To explore the possibility that particular Runx1-dependent protein(s) are maintaining repression, we performed TF binding motif enrichment analysis at these sites. Indeed, Runx motifs were absent from the top 1,500 enriched motifs (Supplemental Table 3), consistent with an indirect mechanism whereby Runx1 acts either by repressing a pioneer activator or by activating a repressor protein. Motif enrichment analysis of Runx1KO-induced ATAC-seq peaks compared to Runx1-dependent ATAC-seq sites revealed significant enrichment for the Zeb repressor motif (Figure 4A; Figure 4-Supplemental Figure 1A). This is consistent with an indirect mechanism in which Runx1 regulates Zeb expression, which in turn actively maintains inaccessible chromatin architecture at multiple sites. Accordingly, we observed that the levels of Zeb1 and Zeb2 mRNA were over 10 fold lower in Runx1KO cells (Figure 4B) and confirmed this independently by real-time quantitative PCR (RT-qPCR) (Figure 4-Supplemental Figure 1D). Using Western blot analysis, we found that the Zeb1 protein is expressed in control cells but not in Runx1KO cells (Figure 4C). Unfortunately, commercially available antibodies to Zeb2 failed to detect it in control cells. Next, we asked if Runx1 was a direct regulator of Zeb1 expression by examining the Zeb1 locus for Runx1 binding. An accessible regulatory region near Zeb1 was bound by Runx1 and Runx2 in control cells (Figure 4D green arrows), but became inaccessible in Runx1KO cells (Figure 4D, red arrows). Similarly, an accessible regulatory region near Zeb2, bound by Runx1 and Runx2 in control cells, became inaccessible in Runx1KO cells (Figure 4E). Additionally, the Zeb1 and Zeb2 TSS, open in control cells, were less accessible in Runx1KO cells (Figure 4-Supplemental Figure 1B), mirroring the dramatic differences in the expression levels of Zeb1 and Zeb2 between control and Runx1KO cells. These data suggest that both Zeb1 and Zeb2 are regulated by Runx1-dependent enhancers in mK4 cells.

**Figure 4:**
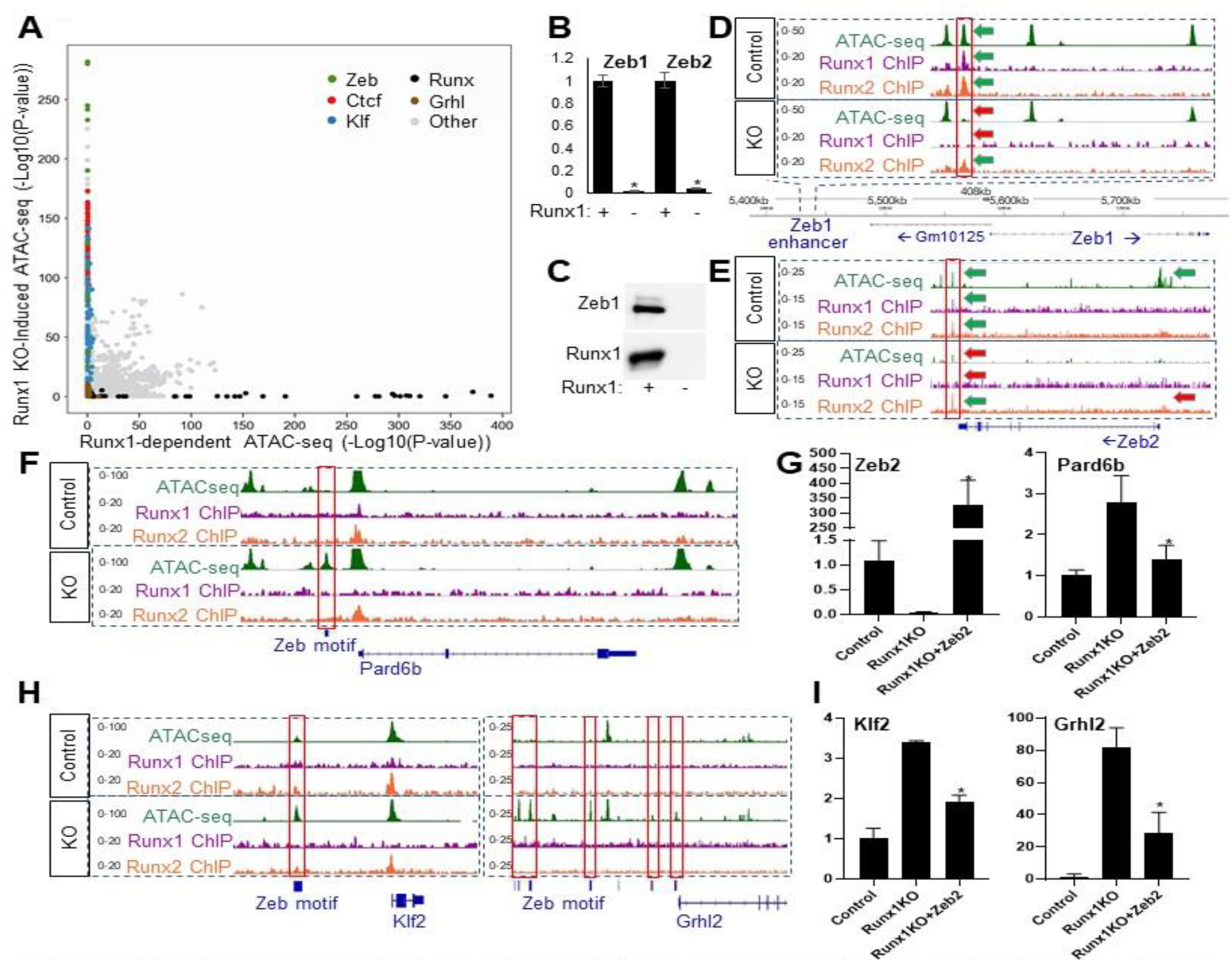
Runx1KO Cells Lack Zeb Repressors, Leading to the Opening of Chromatin. A) Graph displaying p-values of transcription factor motif enrichment in Runx1-dependent versus Runx1-induced ATAC-seq, revealing that Zeb motifs are specifically enriched in Runx1KO-Induced ATAC-seq peaks. Additionally, Ctcf, Klf, and Grhl motifs are enriched in the Runx1KO-induced ATAC-seq peaks, while Runx motifs are enriched in the Runx1-dependent ATAC-seq. Note that this graph has had AP-1 motif enrichment results removed in order to focus on other motif enrichment levels (see Figure 4-Supplemental Figure 1A for all transcription factor motifs). B) RT-qPCR showing that Runx1KO cells lose expression of Zeb1 and Zeb2. C) Western blot showing the absence of the Zeb1 protein in Runx1KO cells. D) Genomic snapshot showing a chromatin region near Zeb1 that is bound by Runx1 and Runx2 and loses chromatin accessibility in Runx1KO cells. E) Genomic snapshot of the Zeb2 locus showing a downstream potential enhancer bound by Runx1 and Runx2 that has decreased chromatin accessibility along with a loss of expression in Runx1KO cells. F) Genomic snapshot of the Zeb target gene Pard6b locus showing a promoter region containing a predicted Zeb binding site that is specifically open in Runx1KO cells. G) RT-qPCR showing that Zeb2 transient transfection of Runx1KO cells induces repression of Pard6b expression down to levels similar to those in control cells after 1 day of selection for transfected cells followed by 2 days of growth in media. H) Genomic snapshots of the Klf2 and Grhl2 loci displaying ATAC-seq regions that are specifically open in Runx1KO cells (outlined in red) that contain predicted Zeb binding sites. I) RT-qPCR confirmation of the upregulation of Klf2 and Grhl2 in Runx1KO cells, which is suppressed by transient transfection of Zeb2, as shown for RT-qPCR of Pard6b (panel G). The * denotes p< 0.05 in Student’s t-test.

In support of the hypothesis that the opening of the chromatin in Runx1KO cells is due to the loss of Zeb repressors, we examined a known Zeb1 repressed target, *Pard6b*, that was upregulated in Runx1KO cells nearly 10-fold based on RNA-seq analysis (Figure 4-Supplemental Data 1C) and RT-qPCR (Figure 4-Supplemental Figure 1D). A predicted Zeb binding site upstream of *Pard6b* is only accessible in Runx1KO cells (Figure 4F).

Thus, we hypothesized that a subset of the genes upregulated in the Runx1KO cells may be the result of losing Runx1-dependent Zeb 1 and Zeb2 expression, and the subsequent derepression of Zeb targets.

To test whether loss of Zeb expression is responsible for the widespread changes in chromatin accessibility and subsequent gains in gene expression seen in Runx1KO cells, we transiently transfected Runx1KO cells with an expression vector driving Zeb2 mRNA levels 330 fold over baseline as determined by RT-qPCR (Figure 4G, left-recall that there is no usable antibody to Zeb2). Next, we tested the expression of *Pard6b* in Zeb2-transfected Runx1KO cells and found that its expression level was significantly reduced (Figure 4G, right). This supports the hypothesis that many of the upregulated genes in Runx1KO cells may be indirectly affected through the loss of Zeb-mediated repression.

Not all of the open chromatin gained in Runx1KO cells contains Zeb sites. Both Klf and Grhl motifs are significantly enriched in the small subset of Runx1KO-induced ATAC-seq fragments that lack Zeb motifs (Figure 4A and Supplemental Table 3). Genomic snapshots of the *Klf2* and *Grhl2* loci reveal Zeb motif-containing chromatin regions that become accessible in Runx1KO cells (Figure 4H), concomitant with increased *Klf2* and *Grhl2* expression, suggesting that Klf2 and Grhl2 are suppressed by Zeb in mK4 cells. As we had observed for Pard6b, transfection of Zeb2 in the Runx1KO cells led to significant downregulation of both *Klf2* and *Grhl2* expression by RT-qPCR (Figure 4I), which supports the interpretation that these genes are normally repressed by Zeb proteins in mK4 cells. The inhibition of Grhl2 expression by Zeb proteins has been shown to play a critical role in reciprocal negative feedback loops between these pathways during the epithelial-mesenchymal transition (Cieply et al., 2013). Combined, these data suggest that upregulation of Klf2 and Grhl2 have functional consequences via opening of chromatin at their bound targets, and support the hypothesis that the upregulation of genes in Runx1KO cells is due in large part to the loss of Zeb repressors, which cascade due to derepression of additional transcriptional activators. Thus, these results reveal how the loss of a single transcription factor can create ripple effects perturbing the entire transcriptional network.

## Discussion

We report herein that Runx1 regulates the chromatin landscape at multiple loci to broadly impact transcription in a mouse kidney cell line. Even though Runx1 has been reported to be both a transcriptional activator and a repressor (Brettingham-Moore et al., 2015; Mevel et al., 2019; Seo et al., 2012), our analyses suggest that Runx1 functions primarily as an activator in mK4 cells despite the expression of Gro/TLE1-3 (Supplemental Table 1). Runx1 does so by maintaining chromatin accessibility at many loci, including near loci encoding the repressors Zeb1 and Zeb2. The Zeb proteins in turn act to maintain inaccessible chromatin at many loci, including several other transcription factors (e.g., Klf2 and Grhl2), resulting in the cascading repression of further downstream indirect targets (e.g., Ovol1). Runx1 deficiency leads to loss of accessibility at regulatory regions of downregulated direct target genes and subsequently to gain in accessibility and upregulation of actively repressed genes, including additional transcriptional activators such as Grhl2 and Klf2 (Figure 5), leading ultimately to widespread genome-wide changes to chromatin accessibility and transcription.

**Figure 5:**
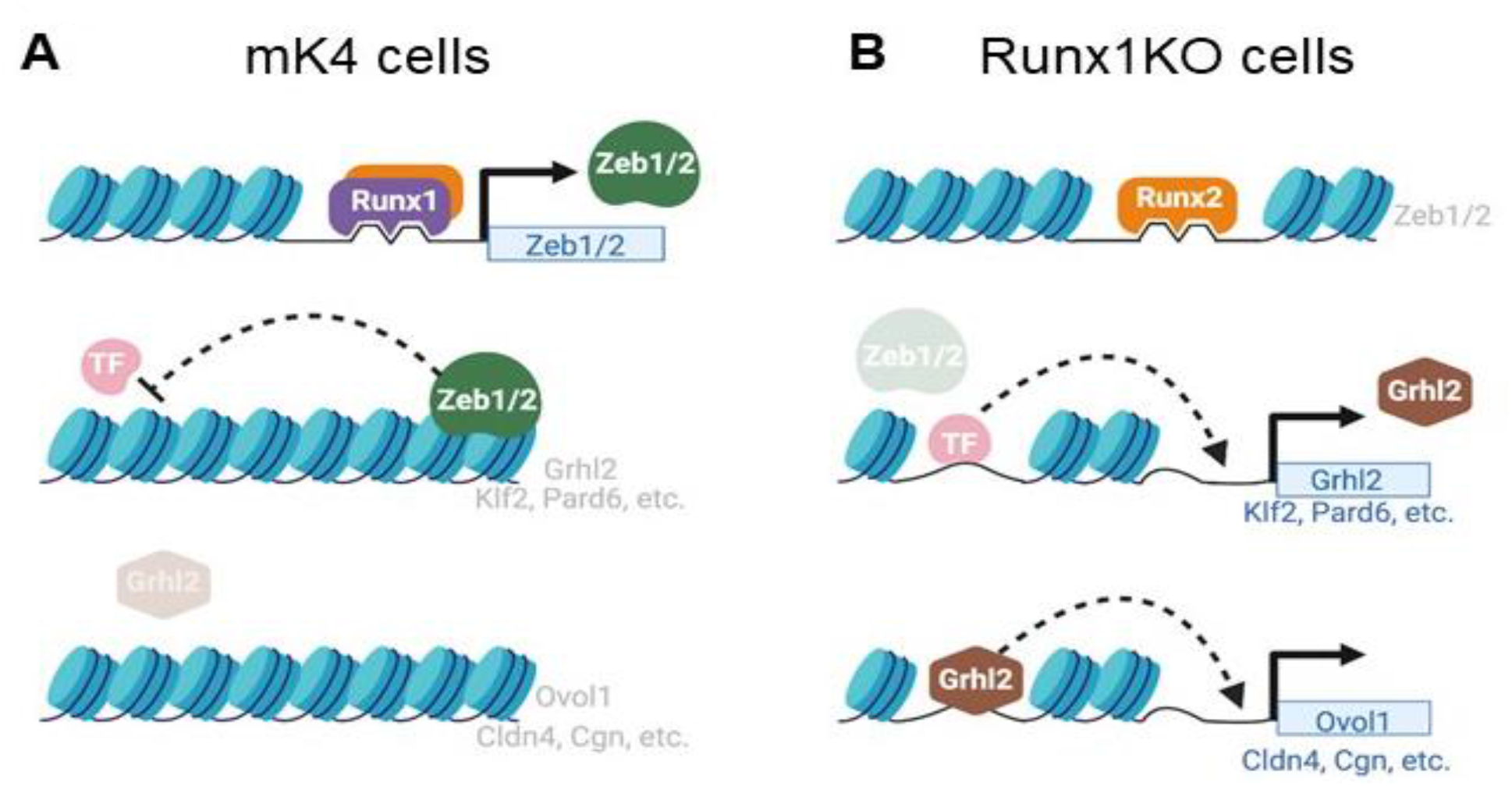
Model of Transcription Factor Network Perturbation by Runx1-Deficiency. A) In control mK4 cells, Runx1 induces the transcriptional repressors Zeb1 and Zeb2 that inhibit other transcriptional activators such as Grhl2, resulting in inhibition of downstream Grhl2 target genes. B) Runx1KO cells lose expression of Zeb1 and Zeb2, which derepresses their targets including Grhl2 and Klf2, which in turn leads to upregulation of their downstream targets such as Ovol1, Cldn4, and Cgn. Figure was created using BioRender.com.

Interestingly, these broad effects on chromatin accessibility and transcription occur despite the presence of Runx2 in these cells. Runx2 remains bound to most of the sites immunoprecipitated by Runx1 but is unable to compensate for Runx1 loss. Despite the conserved ability of both proteins to bind inaccessible chromatin, our results are consistent with Runx1, but not Runx2, acting as a pioneer factor (Lichtinger et al., 2010). The molecular mechanism underlying the specific ability of Runx1 to maintain chromatin accessibility near its target genes remains under investigation.In addition to facilitating transcription by enhancing accessibility, TFs can regulate transcription by recruiting additional factors or by contributing activity to preassembled complexes. In mK4 cells, Runx1 appears to largely act by making chromatin accessible to other TFs and by inducing the expression of repressors to prevent a myriad of other sites from responding to their regulators. The enrichment for down-regulated genes near Runx1-dependent ATAC-seq is greater than that observed for Runx1 ChIP, which suggests that impact on accessibility spreads across chromatin regions beyond the sites directly bound by Runx1. This raises the possibility that Runx1 may play an additional role in facilitating the loading of other TFs through the opening or maintaining of enhancer accessibility, to allow full transcriptional activation of its target genes, as has been described for other transcription factors (van Bakel, 2011). The integration of signaling pathways through control of chromatin accessibility in co-occupied regions, even in the absence of direct interactions, has been shown to be an important regulatory mechanism for other TFs, such as SOX2 and OCT4 (Friman et al., 2019). It will be interesting to determine whether Runx1 plays a generalized master-regulator role controlling the activity of other signaling pathways through the modulation of chromatin accessibility.

The finding that gains in chromatin accessibility in Runx1KO cells largely reflects a loss of Zeb repressor protein activity suggests that Runx1 “repressor” functions may be executed by Zeb1 in many cells and tissues, where Zeb expression requires Runx1. This may reflect an underappreciated and widespread collaboration between these proteins. Accordingly, examination of public functional genomics data revealed that the Runx1-bound Zeb1 enhancer identified in mK4 cells is accessible (DNase hypersensitive) in other tissues including several hemopoietic cell lines (Supplemental Table 4) and is also bound by Runx1 in AML. Further, gaining Zeb expression could be relevant not only to the role of Runx1 in AML but might also contribute to solid organ malignancies, where Zeb proteins are critical inducers of the epithelial-mesenchymal transition (EMT). In agreement with this hypothesis, co-expression of both Runx1 and Zeb2 in circulating tumor cells has been shown to significantly correlate with cancer reoccurrence (Alonso-Alconada et al., 2014). Additionally, Runx1 has been shown to be critical for TGFbeta induced EMT during renal fibrosis both in HK-2 cells and *in vivo* (Zhou et al., 2018). The interplay between the Runx and Zeb calls for a reexamination of whether TGFbeta induction of EMT through increased expression of Zeb proteins (Xu et al., 2009) is mediated through Runx1. Interestingly, the cell-specific dependence of Zeb expression on Runx1 may explain why Runx1 has been suggested to either promote or suppress EMT in different cell lines (Zhou et al., 2018). For example, Runx1 has been shown to inhibit expression of Zeb1 in breast cancer cells, thereby suppressing EMT (Hong et al., 2018; Hong et al., 2017; Li et al., 2019). This raises the possibility that a switch between positive and negative regulation of Zeb proteins by Runx1 could underlie the contrasting reports on its role in promoting or inhibiting EMT. This indirect mechanism by which Runx1 mediates repression through the upregulation of other proteins may be widespread and include additional repressors and activators that could be uncovered by the enrichment for their motifs revealed by experiments such as ATAC-seq in different cellular contexts. Collectively, our data indicate that loss of Runx1 produces widespread genomic and transcriptional changes through a cascade of direct and indirect sequalae involving multiple transcriptional repressors and activators, and reveal key members of this complex network of interacting TFs.

## Materials and Methods

### Tissue culture

The mK4 and Runx1KO cells were grown in DMEM supplemented with 10% FBS, L-glutamine, penicillin/streptomycin, and sodium pyruvate. The cells were transfected using Lipofectamine 2000 (Invitrogen) following the manufacturer’s directions.

### Generation of Runx1KO cells

We used the mK4 cell line as our control cell line, as described in (Valerius et al., 2002). From the control cell line we generated a sub-line that does not express Runx1 through the use of guide RNAs (gRNAs) in the px458 and px459 CRISPR/Cas9 vectors to delete the third exon of Runx1 that contains the start codon utilized in mK4 cells. The targeting sequences were generated with the method and tools described in (Haeussler et al., 2016). These cells were than transfected with Lipofectamine 2000 (Invitrogen) according to the manufacturer’s instructions. The cells underwent selection with puromycin for two days. Subsequent clones were picked approximately a week later using cloning disks and the clones were screened for exon 3 deletion by PCR and for loss of Runx1 protein expression by Western blot.

### Western blot

Confluent control cells or Runx1KO mK4 cells were collected in 100 ul of RIPA-DOC with protease inhibitors plus 100 ul of 2X sample buffer. Protein samples were run on 7% polyacrylamide gels and then transferred to PVDF. Indicated antibodies were applied at 1:1000 dilutions overnight at 4 degrees and then secondary antibodies were used at 1:5000 at room temperature for 1hr. The Western blot signal was detected using Thermofisher Supersignal Femto ECL reagent using a Bio Rad Chemidoc MP Imaging System.

### RNA-seq

mK4 control and Runx1KO cells were cultured in triplicate in standard mK4 media (DMEM plus 10% FBS, 2% L-glutamine, 1% Pen/Strep, and 1% Sodium Pyruvate) on 12 well plates until nearly confluent. Cells were removed from the plate with trypsin that was subsequently inactivated using mK4 conditioned media to prevent a feeding effect from fresh media activating signaling pathways. RNA was collected using Invitrogen’s Purelink RNA Mini kit according to the manufacturer’s directions. RNA-seq on polyA isolated RNA was performed by the CCHMC sequencing core to produce over 20 million reads per sample.

### RT-qPCR

Biological triplicate samples of RNA were converted to cDNA using Superscript II Reverse Transcriptase from Invitrogen following the company’s protocol. The cDNA was diluted to 40 ng/μl, and 5μl of each sample was added to each RT-qPCR reaction that were amplified using iTaq Universal SYBR Green Supermix from Bio-Rad and read on a StepOnePlus Real-Time PCR System from Applied Biosystems. Gene expression levels were normalized to Gapdh and changes were determined relative to control cells, with significance calculated using Student’s t-test.

### ATAC-seq

ATAC-seq experiments were carried out in the control mK4 cells and Runx1KO cells in triplicate. Experiments were performed following the protocol laid out by the Kaestner Lab (Ackermann et al., 2016). The Tn5 used in the experiment was prepared using the method outlined in (Buenrostro et al., 2013). The purification of the library prep was done in accordance with (Corces et al., 2017).

### ChIP-seq

Control and Runx1KO mK4 cells were grown on 10cm plates in triplicate until nearly confluent and removed from the plate using trypsin that was inactivated with conditioned media. Individual cells were counted and 10^6^ cells were used to make the ChIP lysates. Cells were incubated in crosslinking solution (1% formaldehyde, 5 mM HEPES [pH 8.0], 10 mM sodium chloride, 0.1 mM EDTA, and 0.05 mM EGTA in RPMI culture medium with 10% FBS) and placed on a tube rotator at room temperature for 10 min. To stop the crosslinking, glycine was added to a final concentration of 0.125 M and tubes were placed back on the rotator at room temperature for 5 min. Cells were washed twice with ice-cold PBS, resuspended in lysis buffer 1 (50 mM HEPES [pH 8.0], 140 mM NaCl, 1 mM EDTA, 10% glycerol, 0.25% Triton X-100, and 0.5% NP-40), and placed on a tube rotator at 4C for 10 minutes. Nuclei were harvested after centrifugation at 10,000g for 5 min, resuspended in lysis buffer 2 (10 mM Tris-HCl [pH 8.0], 1 mM EDTA, 200 mM NaCl, and 0.5 mM EGTA), and placed on a tube rotator at room temperature for 10 min. Nuclei were collected again by centrifugation at 10,000g for 5 minutes. Protease and phosphatase inhibitors were added to both lysis buffers. Nuclei were then resuspended in the sonication buffer (10 mM Tris [pH 8.0], 1 mM EDTA, and 0.1% SDS). A S220 focused ultrasonicator (COVARIS) was used to shear chromatin (150- to 500-bp fragments) with 10% duty cycle, 175 peak power, and 200 bursts per cycle for 7 min. A portion of the sonicated chromatin was run on an agarose gel to verify fragment sizes. Sheared chromatin was precleared with 20 μl Dynabeads Protein A (Life Technologies) at 4 °C for 1 hr.

Immunoprecipitation of Runx-chromatin complexes was performed with an SX-8X IP-STAR compact automated system (Diagenode). Beads conjugated to antibodies against Runx1 (Rabbit mAb #8529, Cell Signaling) or Runx2 (Rabbit mAb #8486, Cell Signaling) were incubated with precleared chromatin at 4°C for 8 hours. The beads were then washed sequentially with wash buffer 1 (50 mM Tris-HCl [pH 7.5], 150 mM NaCl, 1 mM EDTA, 0.1% SDS, 0.1% NaDOC, and 1% Triton X-100), wash buffer 2 (50 mM Tris-HCl [pH 7.5], 400 mM NaCl, 1 mM EDTA, 0.1% SDS, 0.1% NaDOC, and 1% Triton X-100), wash buffer 3 (2 mM EDTA, 50 mM Tris-HCl [pH 7.5] and 0.2% Sarkosyl Sodium Salt), and wash buffer 4 (10 mM Tris-HCl [pH 7.5], 1 mM EDTA, and 0.2% Triton X-100). Finally, the beads were resuspended in 10 mM Tris-HCl (pH 7.5) and used to prepare libraries via ChIPmentation (Schmidl et al., 2015).

### Processing of functional genomics data

RNA-seq, ATAC-seq, and ChIP-seq reads (in FASTQ format) were first subjected to quality control using FastQC (v0.11.7) (parameter settings: --extract -o output_fastqc -f R1.fastq.gz) (Kalita et al., 2018). Adapter sequences were removed using Trim Galore (v0.4.2) (parameter settings: -o folder -- path_to_cutadapt cutadapt --paired R1.fastq.gz R2.fastq.gz) (Goodwin et al., 2016), a wrapper script that runs cutadapt (v1.9.1) (Bentley et al., 2008) to remove adapter sequences from the reads. The quality-controlled reads were aligned to the reference mouse genome version NCBI37/mm9 using STAR v2.6.1e (Dobin et al., 2013). Duplicate reads were removed using the program sambamba v0.6.8 (parameter settings: -q markdup -r -t 8 trimmed.bam trimmed_dedup.bam) (Tarasov et al., 2015). Gene annotations for RNA-seq analysis were downloaded from the UCSC Table Browser (Karolchik et al., 2004) for the NCBI37/mm9 genome in GTF format.

ATAC-seq and ChIP-seq data were processed using the following steps. Peaks were called using MACS2 v2.1.2 (parameter settings: callpeak -g mm -q 0.01 --broad -t trimmed_dedup.bam -f BAM -n trimmed_dedup_peaks) (Zhang et al., 2008b). Specific ChIP peaks were identified by MACS2 peak calling on the combined replicate reads and removing peaks that overlapped with non-specific background peaks called in the Runx1 ChIP in the Runx1KO cells. Peaks shared across experiments (i.e., peaks shared between replicates or shared between treatments/conditions) were identified as peaks with 50% or greater overlap, using BEDtools v2.27.0 (Quinlan and Hall, 2010). The final peak sets for each condition were obtained by requiring peaks to be present in at least two out of the three biological replicates. When comparing across treatments or conditions, peak overlap between any of the three replicates in either treatment/conditions was considered a shared peak between the treatments/conditions. Final peaks, originally in BED format, were converted to Gene Transfer Format (GTF) format to enable fast counting of reads under the peaks using the program featureCounts v1.6.2 (Rsubread package) (parameter settings: featureCounts --ignoreDup -M -t peak -s 0 -O -T 4 -a common_peaks.gtf -o output_counts.txt trimmed_dedup.bam). The resulting matrix of raw counts was normalized for all experiment types to transcripts per million values (TPMs). TF binding site motif enrichment analysis was performed using the HOMER software package (Heinz et al., 2010), which was modified to use a log base 2 scoring system and the set of mouse motifs contained in build 2.0 of the Cis-BP database (Lambert et al., 2019).

## Acknowledgements

Dr. Eric Brunskill, Hope Rowden, Carmy Forney, Kevin Ernst, and Dr. Xiaoting Chen for critical discussions, technical assistance, organization, and/or computational help in the production of this manuscript. This work was made possible through funding from the following sources: R01 NS099068 to R.K. and M.T.W. and Cincinnati Children’s Hospital “Trustee Award”, “Center for Pediatric Genomics Award” and “CCRF Endowed Scholar Award” to M.T.W.

## Resource availability

All the sequencing data has been deposited at the Gene Expression Omnibus (GEO) and can be located under accession number GSE158093. Additionally, a genome browser session with the data loaded will be made publicly available upon publication of this manuscript. All the cell lines and plasmids used in this manuscript will be distributed upon request.

---

**Figure 1-Supplemental Figure 1.**
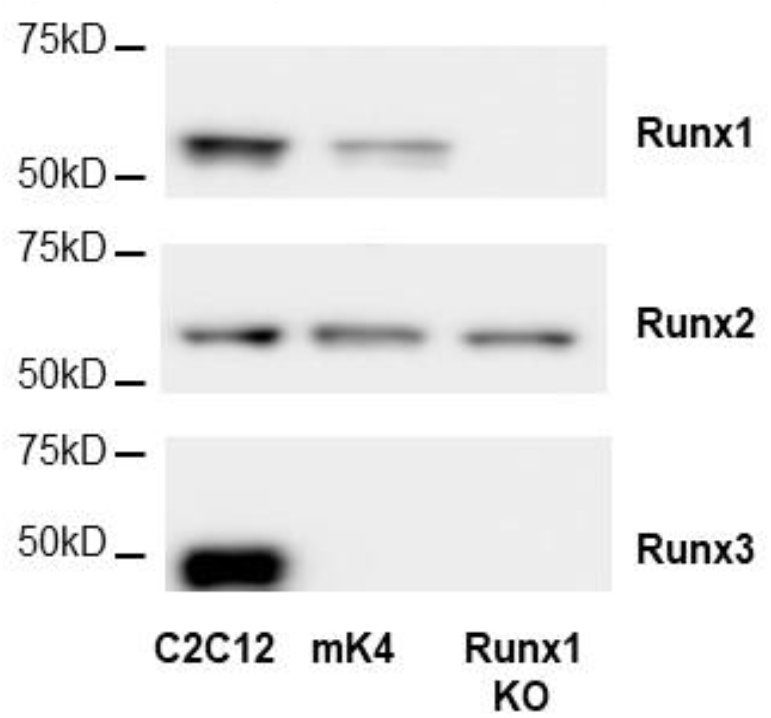
Western blot for Runx1, Runx2, and Runx3 in C2C12, mK4 and mK4-Runx1KO cells showing that all three Runx proteins are expressed in C2C12, Runx1 and Runx2 in mK4 cells and only Runx2 in the mK4-Runx1KO cells.

**Figure 2-Supplemental Figure 1.**
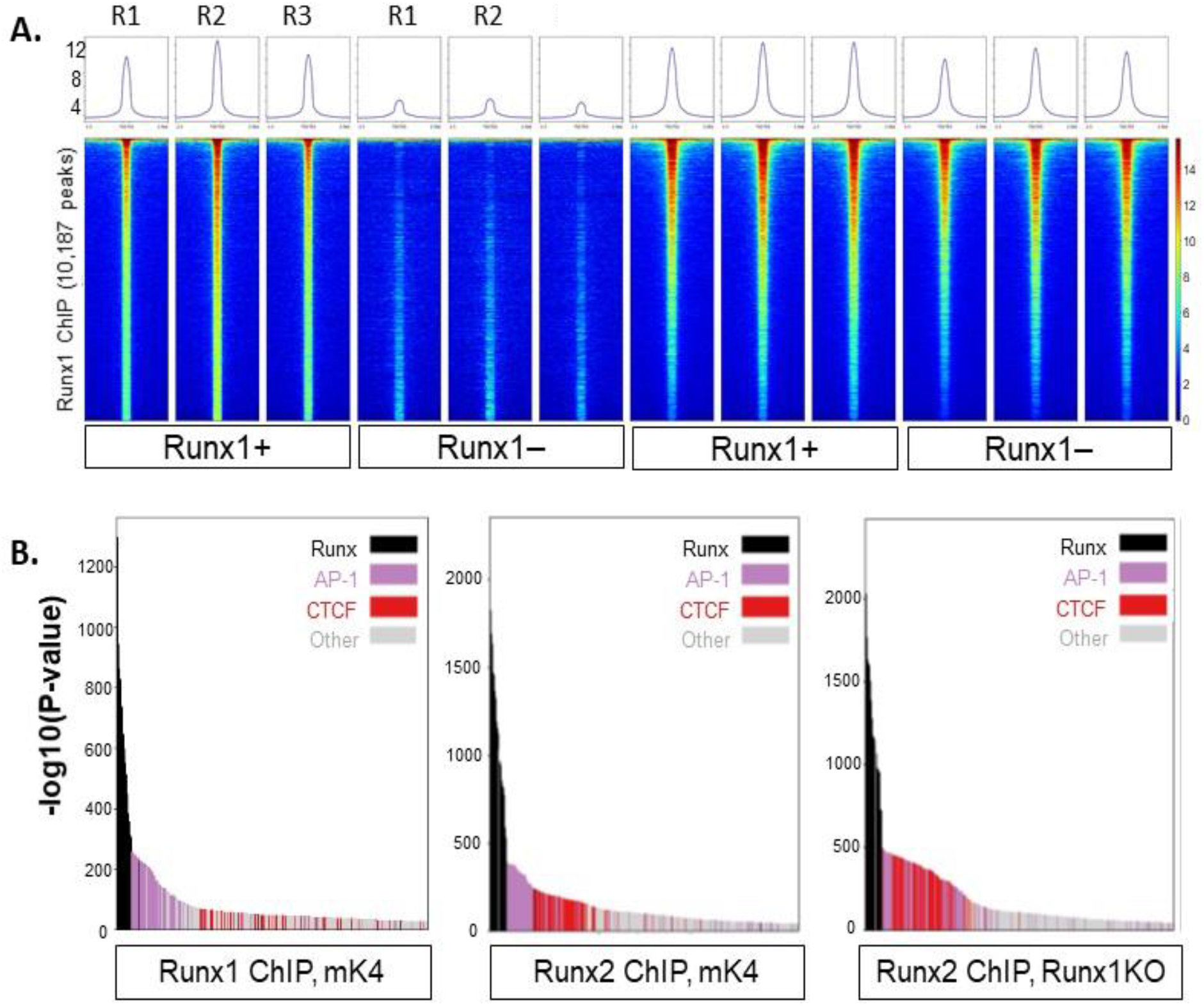
A) Heatmaps of Runx1 and Runx2 ChIP reads mapped to Runx1 ChIP peaks showing the reproducibility of the ChIP replicates. B) Bar graphs of transcription factor motif enrichment −log10 p-values in the Runx1 ChIP in mK4 cells and Runx2 ChIP in mK4 and Runx1KO cells that confirm the strongest enrichment of Runx motifs.

**Figure 3-Supplemental Figure 1.**
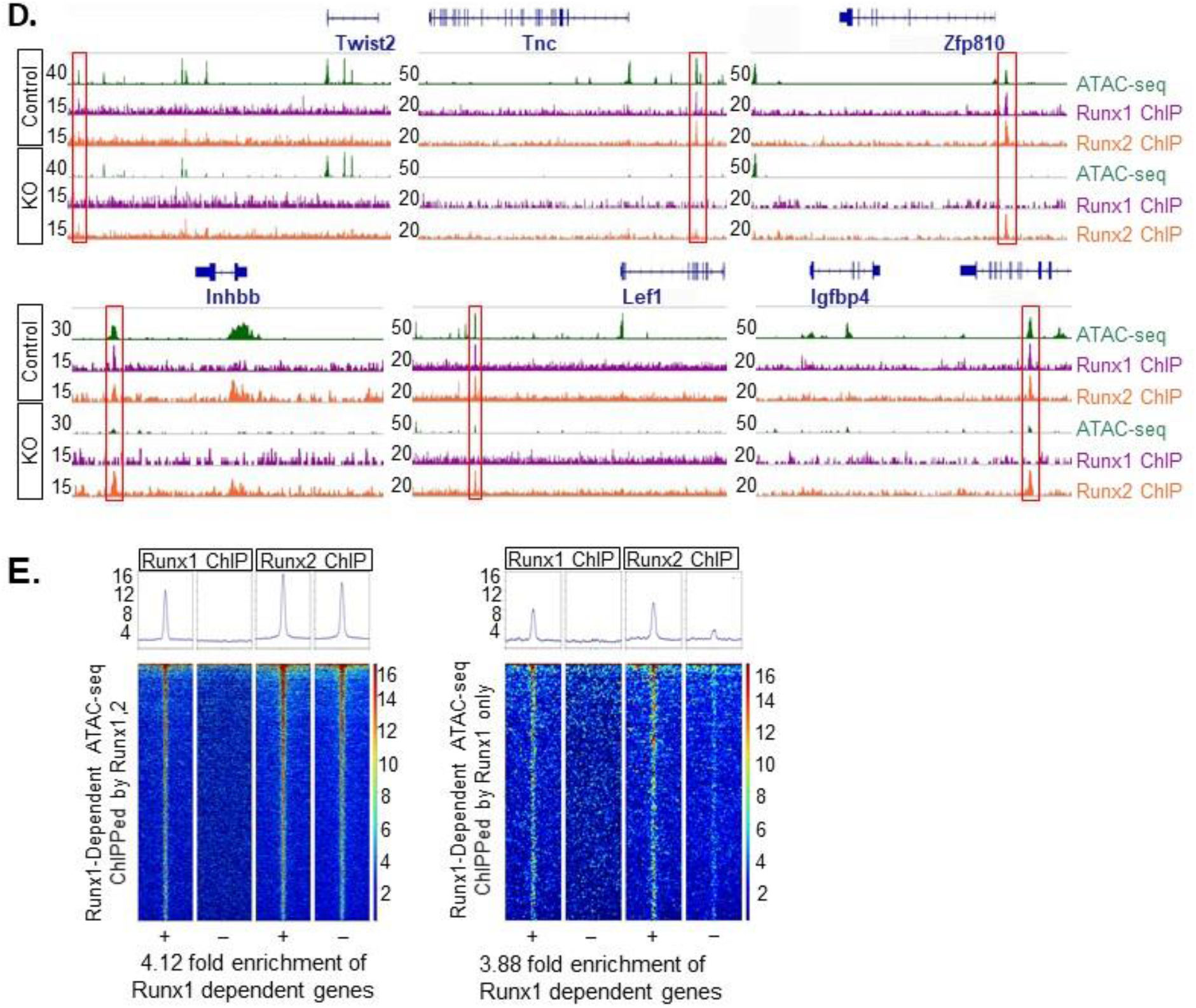
A) Heatmap of Z-scores of individual ATAC-seq samples from mK4 or Runx1KO cells in Runx1-dependent or Runx1KO-induced ATAC-seq peaks showing the reproducibility of the ATAC-seq signal in the replicates and the differences between the mK4 and Runx1KO cells. B) Heatmaps of Runx1 or Runx2 ChIP showing Runx1 binding to both the Runx1-independent and Runx1-dependent ATAC-seq peaks but very little binding to the Runx1KO-induced ATAC-seq peaks. C) Bar graphs of −log10 p-values of motif enrichment in the 3 different classes of ATAC-seq peaks. Runx1-independent ATAC-seq peaks are enriched for AP-1 and Ctcf motifs, Runx1-dependent ATAC-seq enrich for AP-1 and Runx motifs, and Runx1KO-Induced ATAC-seq display enrichment of AP-1, Zeb, and Ctcf motifs. D) Genomic snapshots of Runx1 and Runx2 ChIP and ATAC-seq around the Runx1KO downregulated genes Twist2, Tnc, Zfp810, Inhbb, Lef1, and Igfbp4 that shows Runx1 and Runx2 binding to ATAC-seq peaks that are lost in Runx1KO cells. E) Heatmaps Runx1 or Runx2 ChIP on Runx1-dependent ATAC-seq peaks split into those sites that retain Runx2 binding in Runx1KO cells or sites where Runx2 binding is lost. The Runx1-dependent ATAC-seq that lose Runx2 binding in Runx1KO cells fail to enrich for genes that are down-regulated in Runx1KO cells more than the Runx1-dependent ATAC-seq that retain binding of Runx2.

**Figure 4-Supplemental Figure 1.**
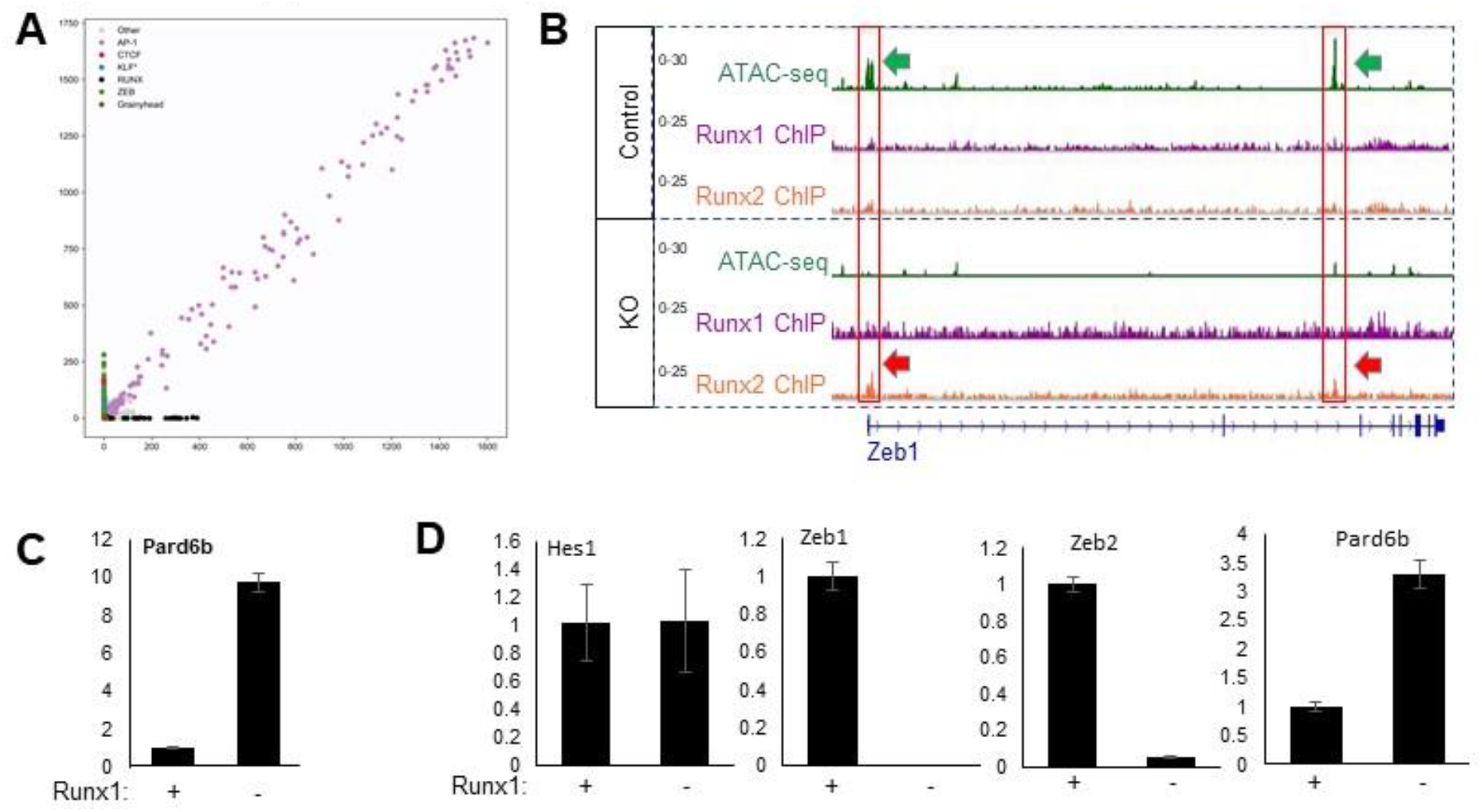
A) Plot showing transcription factor motif enrichment (−log10 p-value) in Runx1-dependent versus Runx1KO-induced ATAC-seq peaks that shows AP-1 motifs are strongly enriched in both but Zeb, Ctcf, Klf, and Grainyhead motifs are specific to Runx1KO-induced ATAC-seq peaks while Runx1 motifs are enriched only in Runx1-dependent ATAC-seq peaks. B) Genomic snapshot of Zeb1 that shows the loss of ATAC-seq signal at the TSS that is lost in Runx1KO cells, which is consistent with the loss of expression in these cells. C) Graph of normalized RNA-seq reads showing the upregulation of Pard6b in Runx1KO cells as expected for a Zeb repressed gene. D) RT-qPCR analysis confirming the dramatic down-regulation of both Zeb1 and Zeb2 and upregulation of Pard6b in Runx1KO cells, but lack of difference for the unrelated gene Hes1.

